# Understanding off-target growth defects introduced to influenza A virus by synonymous recoding

**DOI:** 10.1101/2023.07.17.549395

**Authors:** Colin P Sharp, Beth H Thompson, Blanka Tesla, Dominic Kurian, Peter Simmonds, Paul Digard, Eleanor Gaunt

## Abstract

CpG dinucleotides are under-represented in the genomes of most RNA viruses. Synonymously increasing CpG content of a range of RNA viruses reliably causes replication defects due to the recognition of CpG motifs in RNA by cellular Zinc-finger Antiviral Protein (ZAP). Prior to the discovery of ZAP as a CpG sensor, we described an engineered influenza A virus (IAV) enriched for CpGs in segment 5 that displays the expected replication defects. However, we report here that this CpG-high (‘CpGH’) mutant is not attenuated by ZAP. To understand this, we sought to uncover the alternative attenuation mechanism(s). IAV segment 5 encodes NP, a component of the viral RNA replication complex. Unexpectedly, while CpG enrichment resulted in depleted segment 5 transcript and NP protein abundance, this did not impair viral polymerase activity. A pair of nucleotide changes, introduced as compensatory changes to maintain base frequencies, were instead found to be responsible for the replication defect. These mutations resulted in the encoding of a stretch of eight consecutive adenosines (8A), a phenomenon not seen in natural IAV isolates. Sequencing experiments revealed evidence of viral polymerase slippage at this site, resulting in the production of aberrant peptides and type I interferon induction. When the nucleotides in either of these two positions were restored to wildtype sequence, no viral attenuation was seen, despite the 86 extra CpGs encoded by this virus. Conversely, when these two adenosines were introduced into wildtype virus (thereby introducing the 8A tract), viral attenuation, polymerase slippage, aberrant peptide production and type I interferon induction were apparent. That a single nucleotide change can offset the growth defects in a virus designed to have a formidable barrier to wild-type reversion highlights the importance of understanding the processes underlying viral attenuation. The lessons from this study will inform improved recoding designs in the future.

## INTRODUCTION

To proliferate, viruses must efficiently hijack the host translation machinery to make their own proteins, while subverting cellular antiviral sensors. The evolutionary pressures that these two requirements impart conditions viral genes, manifesting as compositional biases in the genome that become engrained over evolutionary time (1). For example, vertebrate genomes suppress CpG, and consequently, aberrant CpG patterns in viral transcripts may alert the infected cell to the presence of virus. In human cells, this is accomplished through sensing of CpGs in viral RNAs by zinc-finger antiviral protein (ZAP) (2). Viruses with ssDNA, -ssRNA, and some with +ssRNA genomes suppress CpG, thereby mimicking the low genomic CpG content of their vertebrate hosts and evading ZAP-mediated detection (3-5).

Adding CpGs into viral genomes is a now well-characterised attenuation mechanism that has been proposed by ourselves and others to have potential application in live attenuated vaccine design (2,6-13), because adding CpGs limits viral replication but is unlikely to affect antigen conformation. Using influenza A virus (IAV) as a tractable model system in which to test this, we recently reported that increasing the CpG content of segment 1 of IAV resulted in a ZAP-mediated attenuation that caused CpG-high transcript turnover, but not type I interferon induction (14).

Synonymous recoding of viral genomes, including CpG enrichment, is founded on the principle that attenuation is mediated through hundreds of nucleotide changes, negating the potential for reversion to wildtype sequence. However, this does not guarantee that the recoded virus will not revert to wildtype virus fitness. For example, if an RNA structure in a viral genome is required for optimal virus replication, adding CpG dinucleotides would distort that structure, and the virus would be attenuated. It would appear as though the recoded virus was attenuated due to the added CpGs, but in reality the attenuation was mediated by distortion of an RNA structure attributable to one or two nucleotide changes out of the hundreds made. As an historical example of this, a single nucleotide change in the Sabin type 3 live attenuated poliovirus vaccine 5’UTR restored an internal ribosome entry site, improving virus fitness sufficiently to lead to sporadic cases of acute poliomyelitis in some vaccine recipients (15). This highlights the importance of confirming the mechanism when using nucleotide substitutions for viral attenuation.

Prior to the discovery of ZAP as a CpG sensor, we reported that adding CpGs to segment 5 of the A/Puerto Rico/8/1934 (PR8) strain of IAV caused viral attenuation (12). Here, we have tested the sensitivity of this virus to ZAP. Unexpectedly, virus attenuation was not abrogated by ZAP knockout. We find that instead, compensatory mutations made to maintain individual nucleotide frequencies resulted in the introduction of a 8-adenosine (8A) stretch responsible for the attenuated phenotype through IAV polymerase slippage leading to aberrant protein production, associated with an enhanced type I interferon response. Attenuation was alleviated by disruption of the 8A motif, and introduction of these two point mutations into wildtype virus resulted an attenuated phenotype. This study is an important warning, pertinent to the growing interest in using large-scale synonymous virus deoptimisation in vaccinology.

## RESULTS

### CpG enrichment in segment 5 of the IAV genome results in ZAP-independent attenuation

Prior to the discovery of ZAP as the cellular sensor of CpG dinucleotides (2), we had reported the use of CpG enrichment in segment 5 of the PR8 strain of IAV as a successful attenuation approach (12). We therefore tested whether our CpGH IAV was attenuated in ZAP-deficient systems. The virus panel comprised wildtype (WT) PR8, a permuted control with reordered codons but equal dinucleotide composition (CDLR), and CpGH virus, which had 86 CpGs added and further compensatory mutations to retain nucleotide frequencies (**Table 1**). As we previously reported (12), the CpGH virus was significantly attenuated in A549 cells, yielding endpoint titres ∼2log_10_ lower than WT and control viruses. However, in paired A549 ZAP-/- cells, the fitness defect of the CpGH virus was retained (**Fig1A**). Similarly, knockout of other factors in the CpG sensing pathway could not rescue the defective replication of this CpGH virus, either when TRIM25 was knocked out of HEK293 cells (**Fig. 1B**), or KHNYN was knocked out of A549 cells (**Fig. 1C**). The replication defect of the CpGH PR8 virus was therefore not due to ZAP-mediated sensing.

**Figure 1.**
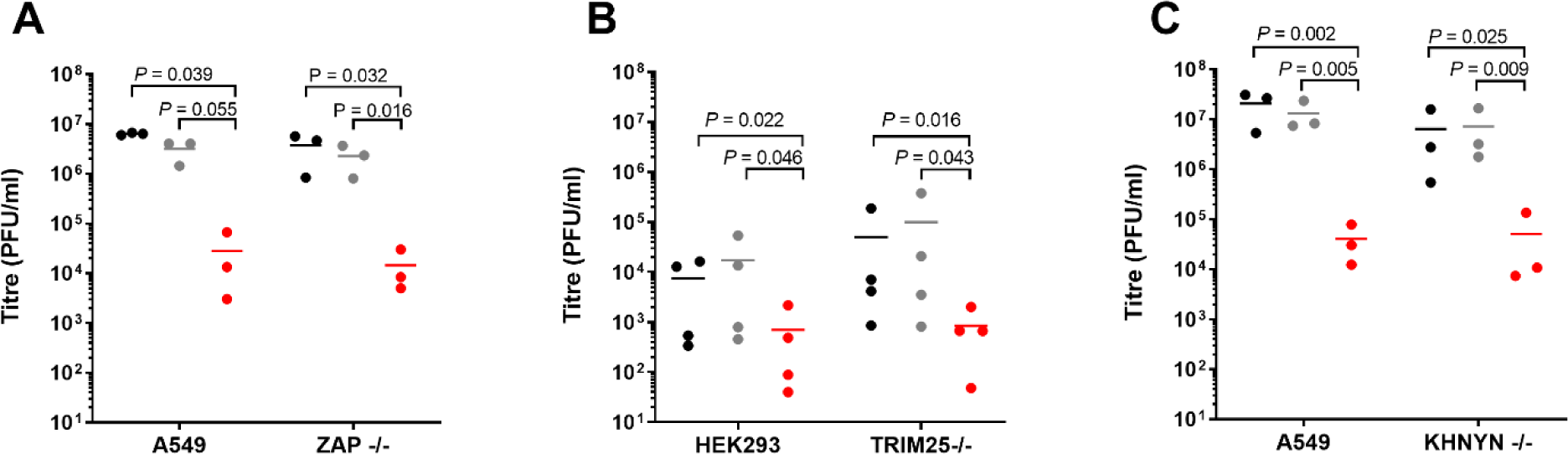
CpG enrichment in segment 5 of the IAV genome results in ZAP-independent attenuation. PR8 WT, CDLR control and CpGH viruses were used to infect permissive cells and counterpart ZAP-pathway knockout cells, and infectious virus production was measured. **A.** WT A549 cell or paired ZAP-/- cell infections. **B.** WT HEK293 cell or paired TRIM25-/- cell infections. **C.** WT A549 or paired KHNYN-/- cell infections.

**Table 1.** Properties of CpG modified PR8 IAVs recoded in segment 5.

| Mutation | Recoded region | No. CpGs | ΔCpG | No. UpAs | Δ UpA |
| --- | --- | --- | --- | --- | --- |
| Wildtype segment 5 | - | 43 | - | 52 | - |
| CDLR | 159-1404 | 43 | - | 52 | - |
| CpGH | 162-1413 | 129 | +86 | 54 | +2 |
| CpGH-A | 628-1413 | 100 | +57 | 53 | +1 |
| CpGH-B | 151-624; 986-1413 | 110 | +67 | 48 | -4 |
| CpGH-C | 151-986 | 91 | +48 | 58 | +6 |
| CpGH-D | 151-297; 783-1413 | 98 | +55 | 49 | -3 |
| CpGH-E | 151-780; 1283-1413 | 94 | +51 | 53 | +1 |

### CpG enrichment in segment 5 of the IAV genome does not impair viral packaging

We considered whether the CpGH virus may be attenuated due to a packaging defect. To test this, RNA was extracted from virus stocks and qPCR was performed to determine the relative amounts of segment 1 and segment 5 RNA in virions. No differences could be found across the panel, in contrast with a known packaging mutant ‘4c6c’, for which more copies of both segment 1 and segment 5 RNA were needed to produce an infectious virus (**Fig. 2A**). RNA from purified virions was inspected visually using urea-PAGE electrophoresis. No differences were observed in band density for any viral segments of any of the viruses in the panel (**Fig. 2B**). CpG enrichment in segment 5 did not impair IAV genome packaging.

**Figure 2.**
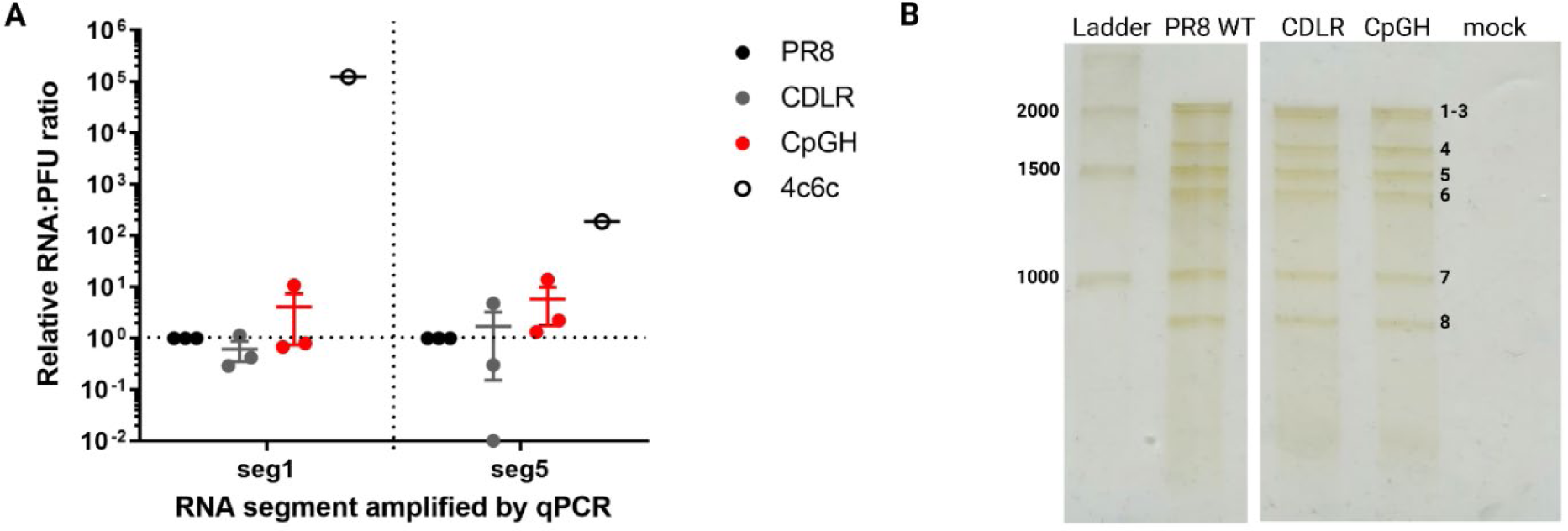
CpG enrichment in segment 5 of the IAV genome does not impair viral packaging. **A.** RNA was extracted from viral stocks for PR8 WT, CDLR control and CpGH viruses and relative copy numbers of segment 1 and segment 5 RNA were quantified by qPCR. A ‘4c6c’ mutant was used as a negative control. **B.** Virus panel virions were purified, then genomic RNA was extracted and individual segments were separated using urea-PAGE.

### CpG enrichment in segment 5 of the IAV genome reduces NP transcript and protein abundance but does not impair IAV polymerase activity

We considered whether CpG enrichment had caused a defect in transcription and/ or translation. Firstly, we tested transcript yield from the CpGH construct in an *in vitro* transcription assay (with RNA produced from plasmids under a T7 promoter); no differences were observed across the panel (**Fig. 3A**). Similarly, no differences in protein levels were detected in *in vitro* translation assays using the same constructs (**Fig. 3B**). Next we examined transcript and protein production during virus infection. A549 cells were infected at high MOI and after a full replication cycle, RNAs were harvested and positive sense viral RNAs were assayed using Northern blotting (**Fig. 3C**). Transcript levels for the CpGH virus were down for both segment 5 (encoding NP) and segment 1 (encoding PB2). This correlated with reduced protein levels (**Fig. 3D**). NP and PB2 both form part of viral ribonucleoproteins (vRNPs), the formation of which is required for viral mRNA production (16,17). We assayed IAV polymerase protein activity by performing viral polymerase reconstitution assays. This involved transfecting plasmids encoding the vRNP proteins (PB2, PB1, PA and NP) under a pol.II promoter, along with a reporter construct in the negative orientation and flanked by IAV UTRs, so that vRNP formation around reporter -RNA is required for translation of the reporter protein. In contrast with the protein production seen during virus infection, in the polymerase assay, equal amounts of PB2 protein were detectable (**Fig. 3E**). While NP protein production was reduced, abundance of the luciferase protein was not reduced. This was confirmed by reading luminescence, as there were no differences in luciferase signal across the panel (**Fig. 3F**), indicating that the viral polymerase proteins were functional. Taken together, these data show that the reduced transcript and protein abundance consequent of CpG enrichment in segment 5 did not impair viral polymerase activity.

**Figure 3.**
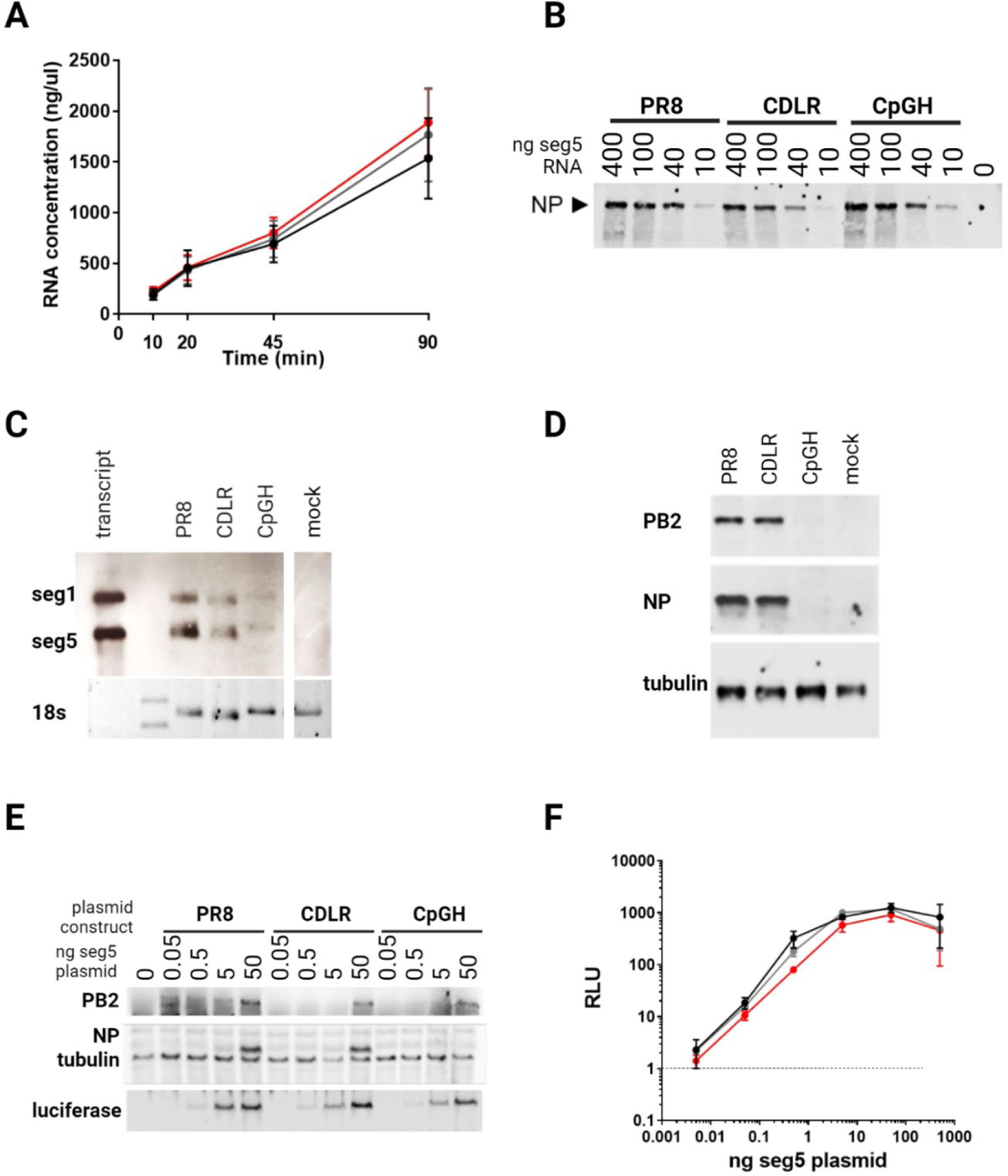
CpG enrichment in segment 5 of the IAV genome reduces NP transcript and protein abundance but does not impair IAV polymerase activity. **A.** To assay transcription efficiency, in vitro transcription assays were performed using the segment 5 plasmids, with RNA quantified at 10, 20, 45 and 90 minutes. **B.** To assay combined transcription and translation in a cell-free system, limiting dilutions of segment 5 plasmids were used in combined transcription and translation assays, and NP protein was detected by western blotting. **C-D.** A549 cells were infected at high MOI for a single viral replication cycle, then RNA abundance was examined by Northern blotting (C) and protein abundance by western blotting (D). **E-F.** Minireplicon assays were performed using limiting dilutions of the segment 5-encoding plasmid for PR8 WT, CDLR and CpGH as a combined measure of transcription and translation in HEK293T cells. E. Cell lysates from the minireplicon assays were probed for viral (PB2 and NP) and reporter (luciferase) proteins by western blotting. F. Luciferase signal was measured by fluorimetry.

### Tiled reversion of the CpGH sequence to wildtype sequence identifies a short region that when restored to wildtype sequence, reconstitutes wildtype virus fitness

To gain insights into the mechanism by which the CpGH virus was attenuated, we sought to identify the recoded region imparting viral attenuation. Segment 5 CpGH was split into 5 fragments, A-E, with A-C forming three sequential fragments covering the full length of the transcript, and D and E mapping across the overlap regions (**Fig. 4A**, **Table 1**). Low MOI infections were performed with this virus panel in A549 cells. CpGH viruses with fragments B, C or E reverted to PR8 WT sequence (CpGH/B.PR8 etc) maintained a replication defect similar to the CpGH virus, indicating that mutations applied in these regions did not contribute to attenuation. However, CpGH/A.PR8 recovered replication fitness to levels similar to the WT PR8 virus (**Fig. 4B**), indicating that the defect in the CpGH virus was imparted by mutation(s) applied in these region. Recovery of the CpGH/D.PR8 virus was evident as it no longer had a titre significantly lower than PR8 WT, although mean titres were almost a log lower than WT PR8. Notably, fragments A and fragments D overlap between nucleotides 298 and 626 (Fig. 4A), raising the hypothesis that mutations in the overlap region may be responsible for the replication defect of CpGH virus.

**Figure 4.**
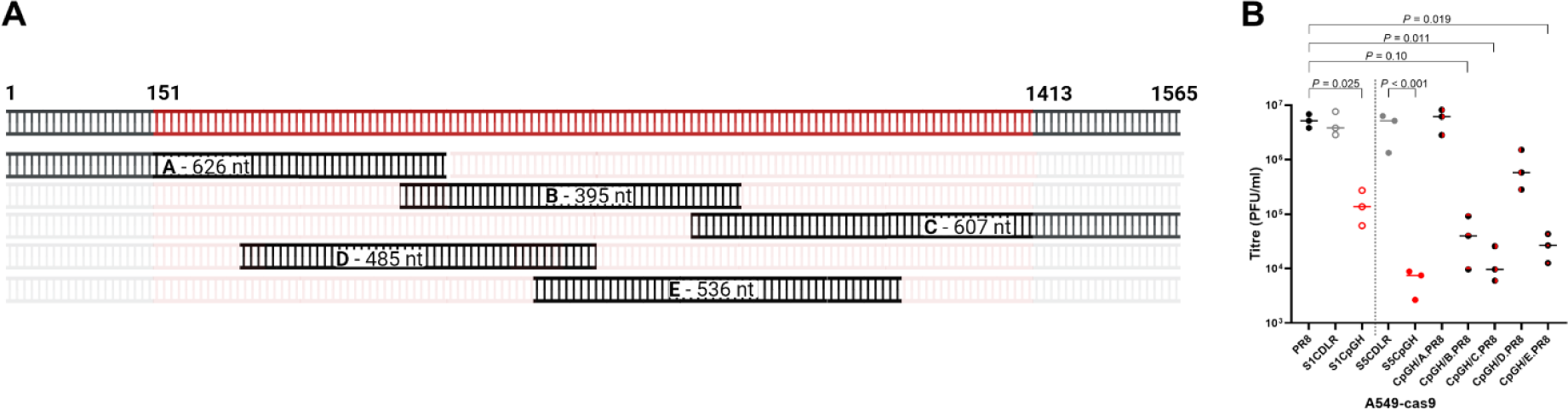
Tiled reversion of the CpGH sequence to wildtype sequence identifies a short region that when restored to wildtype sequence, reconstitutes wildtype virus fitness. **A.** Schematic illustrating reversion of the CpGH sequence in segment 5, achieved using 5 fragments. **B.** Titres of tiled-reversion viruses in A549 cells.

### Serial passage of segment 5 CpGH IAV identifies a reversion mutation within a region mutated to compensate for base frequency changes that falls within an 8-nucleotide stretch of adenosines

To try and identify point mutations responsible for the ZAP-independent attenuation, we performed serial passage experiments. Viruses were passaged at low MOI on A549 cells for 10 passages, and then deep sequencing was performed. Titres did not recover for either egg stock (**Fig. 5A**) or MDCK stock (**Fig. 5B**) CpGH virus. Deep sequencing yielded coverage across the genome between 890-64,000X (**Fig. 5C**). While no nucleotide reversions were seen at sites where CpG dinucleotides had been introduced, a single nucleotide reversion occurred in 98.8% of sequencing reads from both egg virus rescues, at nucleotide position 312. A proximal nucleotide reversion was also seen in one of the MDCK stock rescues, with ∼40% of reads yielding a reversion at position 315 (**Fig. 5D**). Both of these reversions occurred within the overlap region of fragments A and D between nucleotides 298 – 626. To determine when during passage these mutations arose, PCR amplification across this region was performed on RNA extracted from the P1 viral stock and after a further 1 or 2 passages (P2 and P3), and amplicons were deep sequenced. This sequencing showed that the mutation at nucleotide position 312 was present in the P1 viral stock (∼20% of reads), becoming the dominant variant in the next passage and increasing to represent ∼90% of reads by P3 (**Fig. 5E**). This offers a possible explanation as to why the egg stock derived CpGH virus titres were ∼1log below PR8 WT, whereas the difference in MDCK derived virus titres was typically ∼2.5 logs. Visual inspection of nucleotide alignments revealed that the CpGH virus incorporated an 8-adenosine (8A) stretch that included nucleotide positions 312 and 315 (**Fig. 5F**). In over 10,000 human IAV isolate sequences analysed, an adenosine was not seen in positions 312 or 315 (**Fig. 5G**). While sequence reversion at the 8A site was observed, the CpGH virus still displayed a replication defect after reversion at this site (Fig. 5A, B), indicating multi-layered attenuation and consistent with ZAP-mediated CpG sensing.

**Figure 5.**
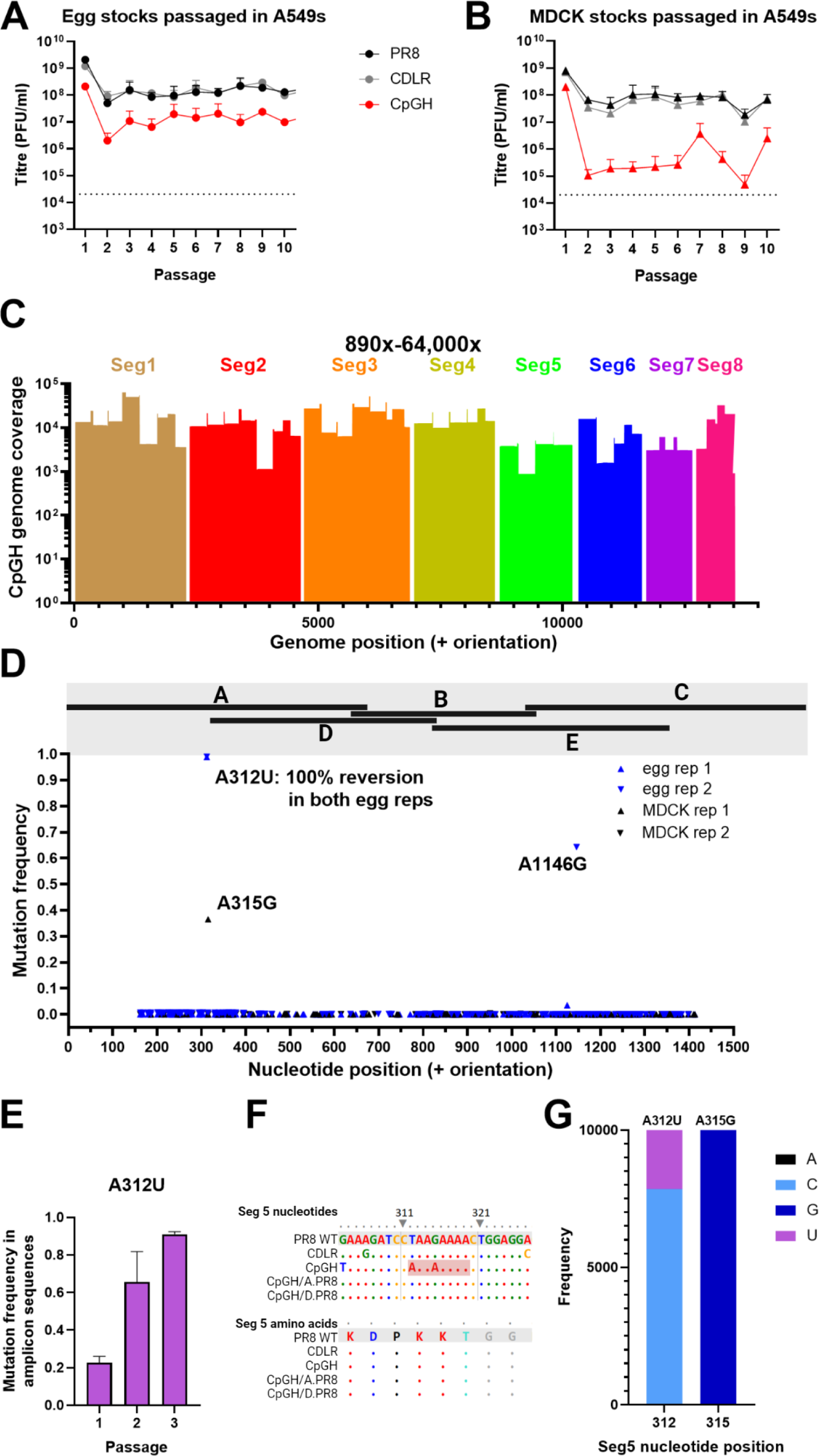
Serial passage of segment 5 CpGH IAV identifies a reversion mutation within a region mutated to compensate for base frequency changes that falls within a 7- nucleotide stretch of adenosines. Virus panel generated from either egg (**A**) or MDCK (**B**) rescue was serially passaged ten times at low MOI in A549 cells, with virus titred after each passage, for four biological repeats (two with starting inoculum of egg rescue and two of MDCK). At passage ten, virus was deep sequenced. **C.** Read depth from deep sequencing of CpGH virus. **D.** Mutations occurring exclusively in the CpGH virus were plotted. None of these mutations occurred at CpG sites. **E.** Egg rescue derived serially passaged viruses were sequenced after 1, 2 and 3 passages; for the CpGH virus, mutation frequency at position 312 is shown. **F.** Reversion mutations at positions 312 and 315 corresponded with an 8-adenosine tract introduced exclusively into the CpGH virus. **G.** Variability of segment 5 nucleotide positions in nature (10,000 human sequence isolates analysed over 5 years between 2018- 2022).

### Single nucleotide reversions at positions 312 and 315 of the CpGH virus restored WT fitness

The tiled reversion data (Fig. 4) and serial passage data (Fig. 5) taken together suggest that the ZAP-independent attenuation of CpGH virus was due to the nucleotide changes at positions 312 and/or 315. Therefore, we made CpGH viruses with these nucleotides reverted singly and in combination, and assessed their replicative fitness in A549 cells. As previously, the CpGH virus was severely defective in comparison with WT and CDLR control viruses (**Fig. 6A**). However, when the 8A tract in CpGH was disrupted by reverting A to U at position 312 (S5CpGH A312U), or the A at position 315 to G (S5 CpGH A315G), the replication defect was lost. Titres were significantly higher than those of CpGH. The same phenotype was observed for the cognate double mutant (S5CpGH A312U/A315G). When these single nucleotide changes were built into PR8, neither PR8 U312A or PR8 G315A were attenuated, but when both mutations were introduced together (PR8 U312A/G315A) to reconstitute the 8A tract, the virus was significantly attenuated compared to WT PR8. Notably, the CpGH virus, which also incorporated 8A but with CpG enrichment built in, was more attenuated than PR8 U312A/G315A, indicating the presence of two different attenuation strategies. Thus, we propose that double-layered attenuation (CpGs enrichment, and the 8A stretch) were acting synergistically.

**Figure 6.**
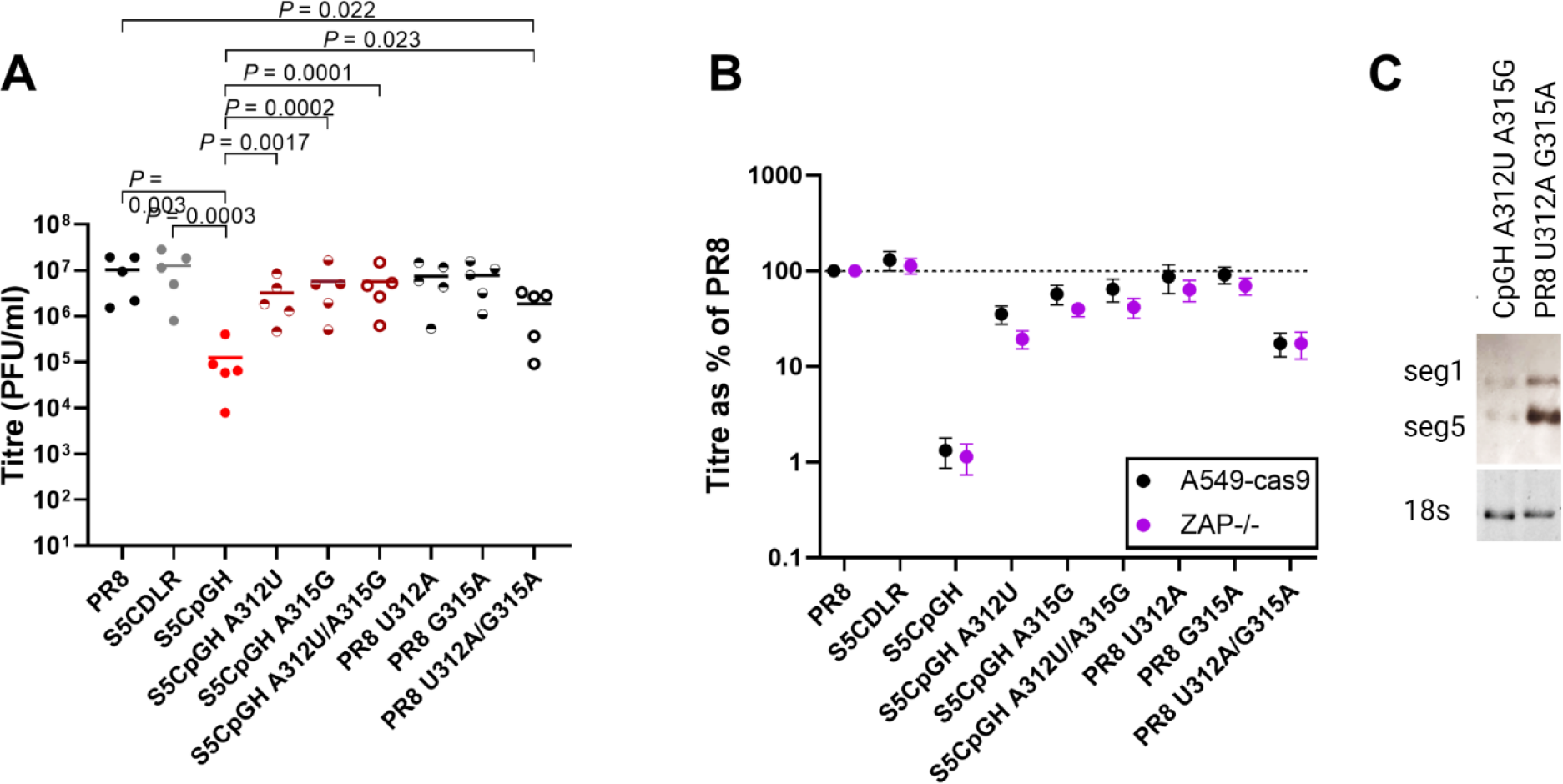
Single nucleotide reversions at positions 312 and 315 of the CpGH virus restored WT fitness. The CpGH virus (‘S5CpGH’) incorporated an 8A tract in the coding sequence. This was removed by reversion to WT sequence at these nucleotide positions through mutations A312U and/or A315G. Conversely, the polyA was introduced into WT PR8 via U312A and/or G315A mutations. **A.** Titre of virus panel grown at low MOI in A549 cells. **B.** To determine whether ZAP sensing was apparent for any mutants, the virus panel was grown in ZAP-/- cells and titres were normalised to PR8 WT titres. **C.** A549 cells were infected with CpGH A312U A315G (no 8A) or PR8 U312A G315A (8A present) at MOI 3 for 10 hours, then RNA was electrophoresed and probed for IAV segments 1 and 5 (positive orientation). Ribosomal RNA (rRNA) serves as loading control.

To determine whether ZAP sensing was evident for any of the CpGH viruses, the same panel was grown in paired A549 ZAP-/- cells. Titres were normalised to PR8 for WT vs ZAP-/- cell lines for direct comparison. No improvement in ZAP-/- cells was observed for any of the viruses in the panel (**Fig. 6B**).

With the presence of 8A determining whether virus was attenuated or not, we sought to test whether its ablation would recover transcript production using Northern blotting as in Fig. 3C. The reduced transcript production seen for the CpGH virus was not ablated by A312U A315G mutations, and could not be imparted upon PR8 by cognate U312A G315A mutations, indicating that reduced transcript levels arose due to CpG enrichment and not 8A introduction (**Fig. 6C**). This was in keeping with our previous observations of ZAP-mediated CpG-high transcript degradation (14), but in segment 5 this alone was evidently insufficient to impair viral replication (Fig. 6A).

### Introduction of the 8A tract into IAV segment 5 resulted in viral polymerase slippage, aberrant protein production, and triggered type I IFN

To determine the mechanism by which the 8A tract introduced into segment 5 caused viral attenuation, we considered the possibility that the viral polymerase was slipping, resulting in aberrant transcript production. To generate the polyA tract at the 3’ end of viral mRNAs, the IAV polymerase is known to slip on genomic polyU (18), although this occurs at the 3’ termini located to the panhandle structure of transcripts rather than in the middle. To test for polymerase slippage, we examined chromatograms of viral transcript sequences when synthesised under different polymerases. Firstly, the CpGH plasmid was transfected into HEK293T cells, and +RNA transcribed from a pol.II promoter was DNAse treated, amplified by RT-PCR across the 312/315 region and sequenced to assess for aberrant transcript production. The sequencing chromatogram indicated no evidence of secondary nucleotide peaks around the 8A tract, indicating that RNA polymerase II did not slip on the 8A sequence (**Fig. 7A, top left panel**). Similarly, when A549 cells were infected with WT PR8 virus, the viral +RNA yielded a clean chromatogram for RNA synthesised by the viral polymerase (Fig. 7A, **top middle panel**). However, when the same cells were infected with CpGH virus whose +RNA contained 8A, downstream of the polyadenosine there was a mixed transcript pool represented in the chromatogram (Fig. 7A, **top right panel**). This indicated that the mixed transcript population most likely arose through viral polymerase slippage. When the A base at nucleotide position 312 was reverted to a U (as in the WT PR8 sequence), evidence of secondary transcript production was absent, indicating that 7 repeated bases was insufficient to generate a slippage event detectable by these methods (Fig 7A, **middle left panel**). Similarly, evidence of slippage was absent from CpGH A315G (containing nucleotide sequence AAAGAAA; **middle middle panel**) and from CpGH A312U A315G (Fig. 7A, **middle right panel**). Finally, the 8A tract was introduced step-wise into the PR8 WT virus; when either U312A or G315A single substitutions were made, no slippage was evident (Fig. 7A **bottom left and middle panels**). However, when both changes were made together, reconstituting the 8A sequence in PR8 +RNA, the mixed transcript population was again evident in the chromatogram (Fig. 7A, **bottom right panel**). Together, these data indicate that a sequence of eight adenosines may give rise to a mixed transcript population (likely via IAV polymerase slippage), and that seven consecutive adenosines is insufficient to yield this effect.

**Figure 7.**
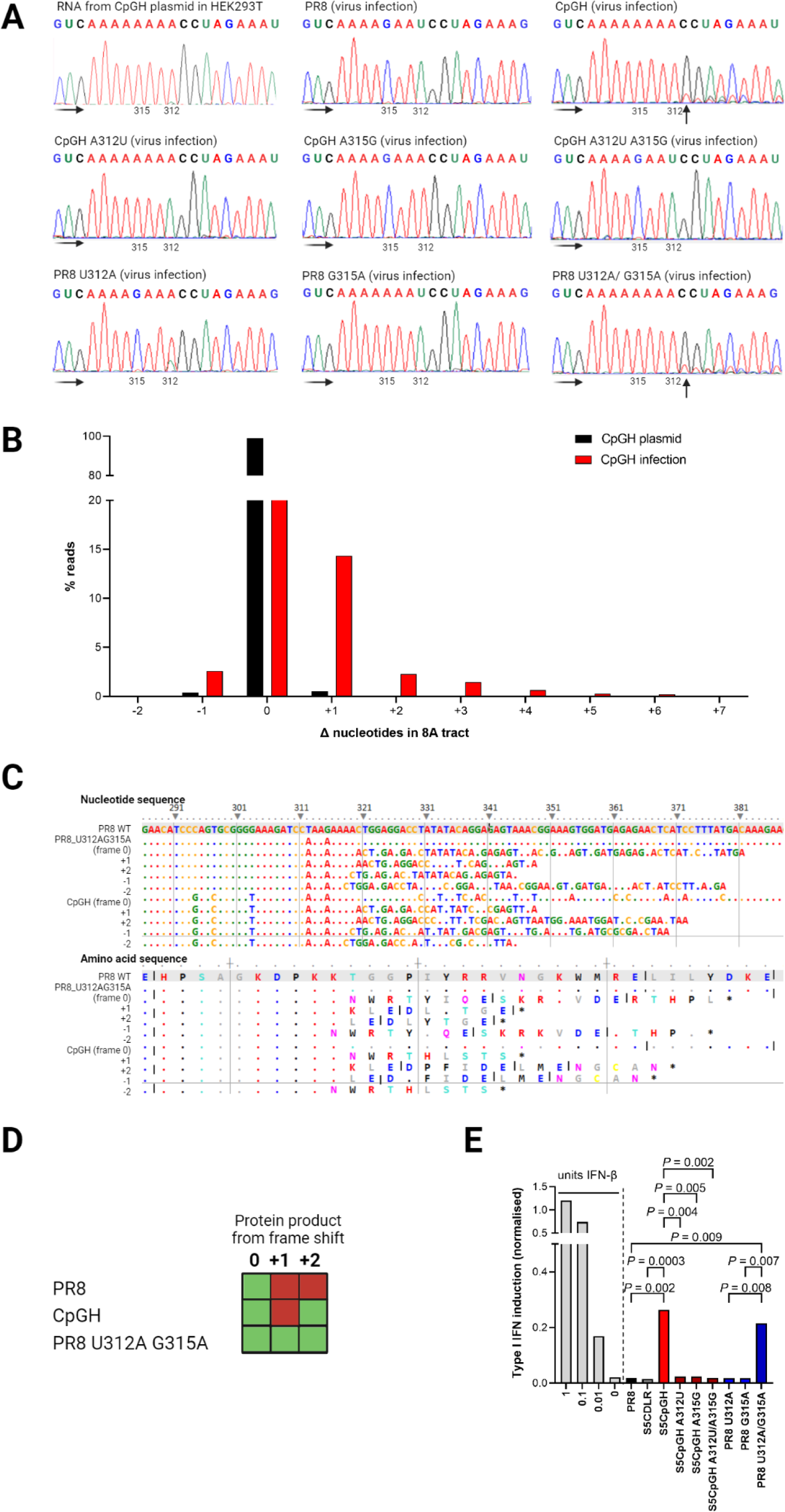
Introduction of the 8A tract into IAV segment 5 resulted in viral polymerase slippage, aberrant protein production, and triggered type I IFN. **A.** WT PR8, CpGH, and cognate viruses with switched nucleotides at positions 312 and 315 were used to infect A549 cells at MOI 3 for 8 hours, after which RNA was extracted, amplified and sequenced. Chromatogram traces were examined for evidence of multiple RNA species generated downstream from the 8A sequence of CpGH virus, encoded at nucleotide positions 312-319. Faded black arrows indicate direction of sequencing read. Solid black arrow indicate sites of polymerase slippage. *Top left*: CpGH plasmid was transfected into HEK293T cells, and +RNA was produced from a pol.II promoter. *Top middle*: PR8 WT virus infection. *Top right*: CpGH virus infection. *Middle left*: CpGH A312U virus (7A tract) infection. *Middle middle*: CpGH A315G virus (4A tract) infection. *Middle right*: CpGH A312U A 315G (4A tract) virus infection. *Bottom left*: PR8 U312A virus (4A tract) infection. *Bottom middle*: PR8 virus G315A (7A tract) infection. *Bottom right*: PR8 U312A G315A virus (8A tract) infection. **B.** CpGH plasmids and infections were deep sequenced and percentage of sequence reads with changes in the length of the 8A tract were calculated. **C.** Top panel – nucleotide alignment of PR8, PR8 G312A U315A, and CpGH sequence surrounding the polyA site, with alignments to show the nucleotide sequence resulting from +1, +2, -1 and -2 polymerase slippage events. The bottom panel shows the resulting peptide species arising from these transcripts. **D.** Presence/ absence of peptide species matrix. Due to differences in m/z ratios for the peptides unique to +1 and +2 frameshifted translations, relative abundance cannot be compared across peptide species. The -1 frameshift followed by gluC digestion resulted in predicted peptides that were not of sufficient length for detection by mass spectrometry. No peptides predicted to have arisen from the -2 frameshift were detected. **E.** Type I interferon competent A549 cells were infected with virus panel for 10 hours at MOI 3, after which time supernatant was harvested, UV treated to inactivate infectious virus and used to treat HEK Blue cells. HEK Blue cells were also treated with IFN standard (light grey bars). IFN induction in these reporter cells was measured by HEK Blue assay. Means of 3 biological repeats are shown.

Deep sequencing of amplicons derived directly from CpGH plasmid versus from virus infection confirmed the increased proportion of sequences containing evidence of polymerase slippage on the 8A site in the context of infection (**Fig. 7B**). This was consistent with the data from Fig. 3, showing that protein but not transcript abundance was reduced in an IAV polymerase-dependent manner.

To assess the consequences of IAV polymerase slippage at 8A on protein production, cell lysates from PR8 WT, CpGH and PR8 U312A G315A infections were analysed using mass spectrometry to determine whether aberrant proteins were produced. Infected cells were treated with MG132 protease inhibitor to prevent their turnover. Sequencing data (Fig. 7B) indicated that frame shifts were most likely to occur in a +1, +2 or -1 orientation, and so the predicted protein translations resulting from such shifts (**Fig. 7C**) were searched for. Due to the 8A stretch introducing multiple lysines, this meant that the standard approach of tryptic cleavage would not allow us to distinguish between +1 and -2, or +2 and -1 frame shift events, and so gluC protease which cleaves downstream of glutamic acid was used instead. This meant that we were unable to detect peptides associated with a -1 frameshift, due to predicted peptides being too short to read. We therefore examined for the presence of peptides associated with canonical, +1, +2 and -2 reading frames (**Fig. 7D**). Due to differences in m/z ratios for the peptides unique to +1 and +2 frameshifted translations, relative abundance cannot be compared across peptide species. For WT PR8, only peptides in the canonical reading frame were identified. For CpGH virus, canonical and +2 frame peptides were identified, but +1 peptides were not. For PR8 U312A G315A, canonical, +1 and +2 frame peptides were identified. No -2 frame peptides were identified in any samples. The detection of +2 frame peptides for CpGH virus suggests that +1 peptides were likely present but of too low abundance or required higher sensitivity detection methods for their identification. Thus, the mass spectrometry confirmed that IAV polymerase slippage resulted in production of aberrant proteins.

We considered the possibility that aberrant transcriptional and translational events may result in the induction of type I interferon, which was investigated using HEK Blue assays. As expected due to the potent type I interferon blocking activity of NS1 (19), PR8 WT and CDLR control did not induce interferon above baseline (**Fig. 7E**). Conversely, 8A-containing CpGH virus significantly induced interferon, and this induction was abrogated in the CpGH A312U, A315G and double mutants. In support of this, PR8 A312U A315G (8A-encoding), but not the single mutants, also induced interferon. Together this indicated that production of the aberrant peptide was required for type I interferon induction.

### Seg1-CpGH virus attenuation is not augmented by incorporation of seg5-CpGH

We sought to determine whether the ZAP-sensitivity of our seg1-CpGH virus could be augmented by incorporating further CpG enrichment into segment 5 of the PR8 genome. To test this, we engineered viruses that were either permuted CDLR controls (no change in CpG frequency) in both segments 1 and 5, or were CpG-enriched in both segments, with segment 5 also incorporating A312U/A315G reversions to negate the risk of polymerase slippage. Firstly, these viruses were grown in WT A549 cells. The seg1-CpGH virus was attenuated as expected, consistent with our previous work (**Fig. 8A**). The seg1/ seg5 CDLR virus yielded titres similar to WT PR8, indicating that synonymously recoding both segments did not impair viral packaging or replication. When CpGs were added into both segments 1 and segment 5, this virus was only attenuated to the extent of seg1-CpGH alone. This indicated that synergistic attenuation by incorporating more CpGs was not achieved. As expected, both CpGH viruses (single and double segment mutants) replicated to WT titres in ZAP-/- cells (**Fig. 8B**). Finally, titres were normalised to PR8 in A549 and paired ZAP-/- cells to assess the extent of recovery in ZAP-/- cells (**Fig. 8C**). Recovery of single and double segment CpGH viruses was similar.

**Figure 8.**
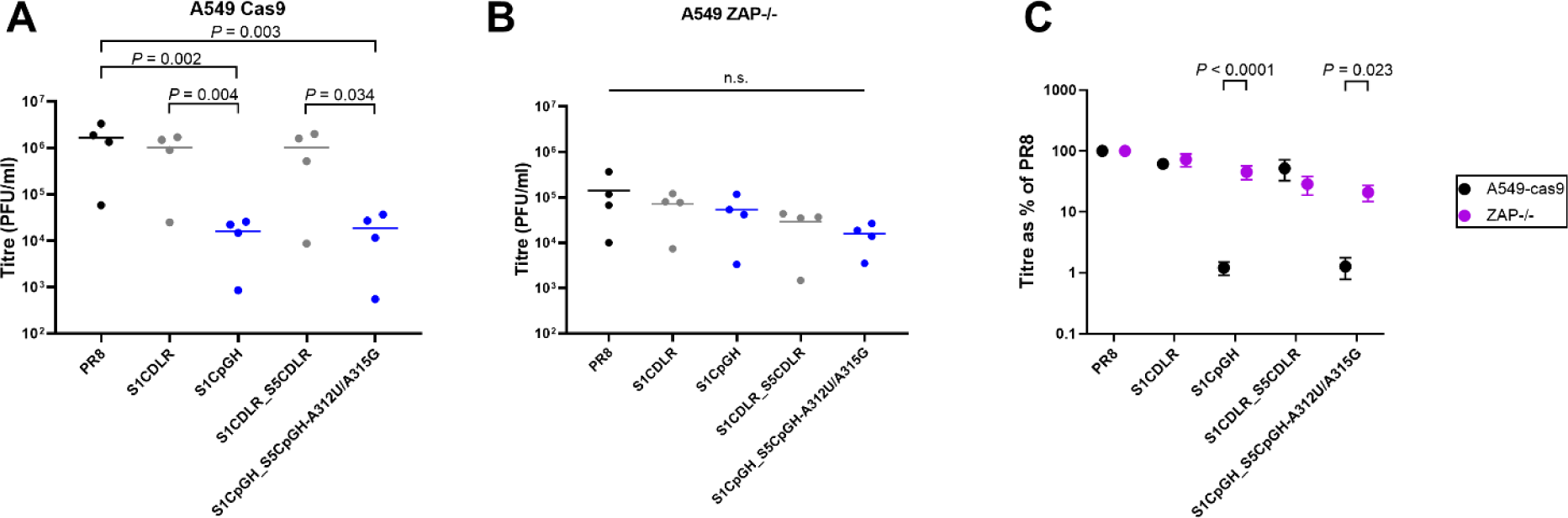
Adding CpGs to segment 5 of a seg1-CpGH virus does not augment the attenuation phenotype. Our previously published seg1-CpGH virus, known to be ZAP-sensitive, was combined with segment 5 mutants to engineer 6:2 virus rescues with segments 1 and 5 both recoded. **A.** Virus panel was used to infect A549 cells expressing cas9. **B.** Virus panel was used to infect ZAP-/- cells. **C.** Titres were normalised to WT PR8 to assess recovery of fitness in ZAP-/- cells.

## DISCUSSION

CpG enrichment has gained traction as an attractive model for the development or augmentation of live attenuated vaccines (13,14). Here, we argue that understanding the mechanism of attenuation imparted by synonymous recoding is imperative as a safety feature of this technology. When we added 86 CpGs to the IAV genome (12), the virus was attenuated and phenotypically characteristic of a virus that was being sensed by ZAP. However, through molecular investigations we have now determined that the replication defects observed primarily arose through an unrelated mechanism requiring a single nucleotide change for fitness restoration. This is therefore a possible scenario during live attenuated vaccine design.

When we added 126 CpGs to segment 1 of the same PR8 IAV, a ZAP-mediated attenuation was imparted, and ∼80 CpGs was sufficient to mediate attenuation. Here, the addition of 86 CpGs without also adding the 8A tract (S5CpGH A312U/A315G) imparted a modest attenuation, evident in Fig. 4 and the maintained replication defect after serial passage (Fig. 5A, B), but evading statistical significance in Fig. 6. The bigger impact of adding CpGs into segment 1 could simply be attributable to the different expression levels of the two genes (20). NP encoded on segment 5 is produced in high abundance (21), and so it is possible that some degradation of viral transcripts is tolerable. In support of this, we saw that while NP production was dramatically impaired during minireplicon assays, this did not impact polymerase activity. This highlights another previously unreported but important consideration for CpG-based recoding designs, as more robust phenotypes may be evident when targeting process-limiting transcripts (14).

Alternatively, RNA structure around CpG sites may determine transcript fate. Binding of CpG motifs by ZAP requires the surrounding RNA to be single stranded (22). TRIM25 is required for ZAP’s RNA-degrading activity (23,24), but while TRIM25 is known to bind RNA, no specific motif or structure has been identified for TRIM25 recruitment (25,26). It is unknown whether there is a minimum threshold of ZAP binding to designate transcripts for degradation. CpG enrichment experiments have so far, by necessity, taken crude approaches of adding CpGs to excessive abundance, whereas it may be possible to strategically add fewer CpGs in regions of single-strandedness (and without introducing RNA structure) to deliver the same attenuation phenotype.

It was unexpected that combining CpG enrichment in segment 1 and segment 5 together did not augment attenuation. We had expected that if cells were primed to degrade segment 1 CpGH transcripts as we previously showed (14), there would be a knock-on effect of enhanced degradation of seg5-CpGH transcripts also that would result in enhanced attenuation. Possibly, the RNA degradation pathway accessed via CpG enrichment was already saturated. This hints at a possible limitation of CpG enrichment as a means for viral attenuation that has not previously been reported.

The finding that the IAV polymerase was slipping at the 8A site was surprising, as such slippage has only been reported on the polyU tract at the 5’ end of negative sense RNA (18), to produce a polyadenylation signal on viral mRNAs. Here, we provide evidence of polymerase slippage in the middle of a transcript. Our assays do not definitively determine whether the polymerase slips on the 8A of positive sense RNA, or the 8U of negative sense, and it was shown that if polyU was replaced with polyA, IAV polymerase would also slip on the polyU (18). Slippage on polyU at the segment terminus requires a tract of six consecutive uridines, in contrast with our finding that 8-nucleotide repeat but not a 7-nucleotide repeat induced slippage. It could be that the polymerase slips on polyA alone, or the differences may be attributable to the different stoichiometry in the middle of the segment compared with the panhandle structure in which the native polyU sequence resides.

With the advent of mRNA vaccine technology, the future of live attenuated vaccines is increasingly uncertain, including for influenza (27,28). Cold-adapted, live attenuated influenza vaccines are typically inoculated in children using nasal sprays (29-32), thereby avoiding the use of needles; a feature that is unlikely to be achieved by mRNA vaccines. Furthermore, synonymous recoding may have applications in both live-attenuated and in mRNA influenza vaccinology in the future (33-35); despite their relative infancy, the SARS-CoV-2 vaccines are codon-optimised (36). Here, we highlight the potential pitfalls of recoding-mediated attenuation, with important implications for vaccine safety.

## METHODS

### Cells

A549 (human adenocarcinoma), Madin-Darby canine kidney (MDCK), and human embryonic kidney (HEK) 293T or 293 cells were cultured in Dulbecco’s modified essential medium (DMEM) (Sigma) with 10% foetal calf serum (FCS) (Thermo Fisher) and 1% penicillin/ streptomycin (Thermo Fisher) (growth medium), and were passaged twice weekly. A549 ZAP-/- cells were a gift from Prof. Sam Wilson (37). HEK 293 TRIM25-/- cells were a gift from Prof. Gracjan Michlewski (26). A549 KHNYN-/- cells were a gift from Dr Chad Swanson (38). The knockout status of these cell lines was previously verified by our lab (14). Cells were checked monthly for mycoplasma contamination using Lonza MycoAlert kit.

### Plasmids and plasmid mutagenesis

IAV reverse genetics plasmids for the A/Puerto Rico/8/1934 (PR8) virus were derived from the UK National Institute of Biological Standards & Control strain (39). The CpGH virus, with CpGs added into segment 5 of the PR8 strain, has been previously described (12). To generate a panel of reversion mutants with fragments of the CpGH region reverted to wildtype sequence, PCR and Gibson assembly were used. For PCR reactions, fragments of either the WT PR8 segment 5 plasmid, or seg5-CpGH plasmid, were amplified (as was the plasmid backbone, which is the same for both constructs; primers are listed in Table S1A). The fragments generated are summarised in Fig. 1A and Table 1. Generated fragments and backbone were added in 1:1 ratios directly to the Gibson assembly reaction mix equivalent to half the total reaction volume, and reactions were carried out in accordance with the manufacturer’s instructions (NEB, M5510A) to generate new plasmids, confirmed by agarose gel electrophoresis. These were heat-shocked into DH5α chemically competent *Escherichia coli*, and grown on agar plates containing 50 µg/ml ampicillin selection. Picked colonies were sequenced using colony PCR, and plasmids containing the correct insert sequences were amplified using Qiagen midi-prep kit, then checked again by Sanger sequencing.

Introduction of single nucleotide changes in segment 5, corresponding to nucleotide positions 312 and 315, was achieved using QuikChange II site directed mutagenesis kit (Agilent) according to manufacturer’s instructions. Primers are tabulated (**Table 2**).

**Table 2.**
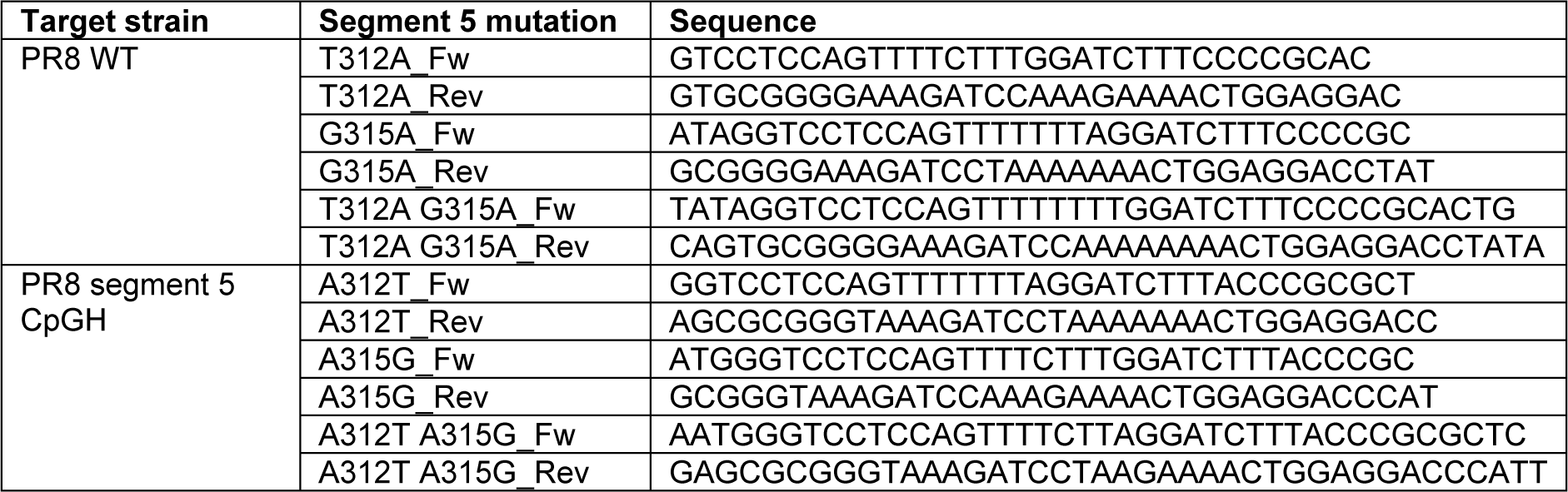
Site directed mutagenesis primers targeting nucleotide positions 312 and 315 in PR8 segment 5

Full segment sequences are summarised (Table 1).

### Virus rescues

Virus rescues were performed as previously described (12,40,41). Growth medium on HEK293T cells at 90% confluency in 6 well plates was replaced with Opti-MEM reduced serum medium (Thermo Fisher) and cells in each well were transfected with 250ng each of 8 pDUAL reverse genetics plasmids, one per segment of viral genome, in combination with 4 µl Lipofectamine 2000 (Thermo Fisher). Mocks were transfected with 7 plasmids, with segment 5 plasmid omitted. The next day, Opti-MEM was replaced with DMEM containing 1 µg/ml tosyl phenylalanyl chlormethyl ketone (TPCK)-treated trypsin and 0.14% bovine serum albumin (w/v) (Sigma) (viral growth medium). After 48 hours, supernatants containing viruses were collected and inoculated into the allantoic fluid of embryonated hens’ eggs at 10 days post-fertilisation (100 µl/ egg). To generate a control virus defective in packaging, a virus with several amino acid changes in segments 4 and 6, encoding the surface glycoproteins, referred to as ‘4c6c’, was propagated in MDCK cells rather than eggs to minimise the risk of reversion (Gog 2007 NAR; eNP).

### Virus titrations

The amount of infectious virus in virus stocks and infection supernatants was quantified using plaque assays. 300 µl volumes of ten-fold serial dilutions were inoculated onto confluent MDCK cells in 12-well plates. After ∼1 hour, cells were overlaid with viral growth medium diluted 1:1 in 2.4% cellulose (Sigma). Plaque assays were incubated for 48 hours, then 1ml/ well 10% neutral buffered formalin was added to fix the cells for at least 20 minutes. Overlay was then discarded and cells were stained using toluidine blue dye (0.1%) (Sigma).

### Virus infections

For viral growth assays and serial passage experiments, cells were infected at low MOI (0.01) and infected for 48 hours in viral growth medium. For interferon induction, replication kinetics and mass spectrometry experiments, high MOI (3) infections were performed for 8 hours in serum-free medium. For all infections, virus was incubated on cells for ∼1 hour, after which time (considered to be 1 hour post infection) cells were washed and the relevant medium was added.

### Virus purification and Urea-PAGE gel electrophoresis

10^10^ PFU of egg-derived virus stocks were semi-purified by pelleting through a 25% sucrose cushion (100 mM NaCl, 10 mM Tris-HCl pH 7.0, 1 mM EDTA) using centrifugation in a Beckman Coulter Max-E ultracentrifuge at 280,000 *g* for 90 minutes at 4C with a SW32Ti rotor. Pelleted virions were resuspended into 350 µl RLT buffer and RNAs were extracted with an RNEasy mini extraction kit (Qiagen). RNAs were loaded onto homemade 5% urea polyacrylamide gels in 1X Tris-borate-ETA (TBE) buffer (89 nM Tris-borate, 2 mM EDTA pH8.3) and separated by electrophoresis at 120 V for 6 hours. RNA as visualised using Bio-Rad Silver Stain Plus Kit and a Samsung Xpress C480FW scanner.

### *In vitro* transcription assay

To generate DNA input under a T7 promoter suitable for use in cell-free transcription assays, full length segment 5 was amplified by PCR using primers with a T7 promoter sequence added to the 5’ end of the forward primer (Table S1A). Q5 DNA polymerase kit (NEB) was used to make amplicons, with 25 ng plasmid and an initial incubation of 95°C for 5 minutes, followed by 30 cycles of 95°C for 30 seconds, 50°C for 30 seconds and 72°C for 2 minutes, then a 72°C for 5 minute final extension. PCR products were run on an agarose gel, then bands of the expected size were excised and gel purified using Qiagen MinElute Gel Extraction Kit. DNA yield was quantified using Qubit dsDNA BR Assay (Thermo Fisher). 40 ng amplicon was used as template in MEGAscrpt T7 transcription assays (Thermo Fisher) with half-reaction volumes, for 10, 20, 45 or 90 minutes. Reactions were terminated and RNA was quantified using Qubit RNA HS Assay (Thermo Fisher).

### *In vitro* translation assay

RNA transcripts generated as above were used as input templates for cell-free translation assays using Promega Rabbit Reticulocyte Lysate kit with one fifth volume reactions. Transcend Biotin-Lysyl-tRNA (Promega) was incorporated in reactions so that reaction products could be identified using western blotting as above, except that membranes were blocked using 5% BSA/TBS for 60 minutes, then incubated with IRDye 800CW Streptavidin (LICOR) for a further 60 minutes, and then imaged using LICOR Odyssesy Fc imaging system.

### Western blotting

Cell lysates from each well of a 24-well plate were harvested in 100 µl Laemmli buffer (2X) and boiled for ten minutes. Samples were cooled and 5 µl/ well was loaded into 10% polyacrylamide precast gels (Bio-Rad) and SDS-PAGE was performed. Resolved proteins were wet-transferred onto nitrocellulose memrbanes (Fisher Scientific) at 100V for 90 minutes. Membranes were blocked for 30 minutes with 5% skimmed milk powder diluted in PBS (w/v) and 0.1% Tween-20. Membranes were probes with NP, PB2 (42), firefly luciferase (EPR17790, Abcam) (all 1:1000) and β-tubulin (clone YL1/2, Bio-Rad; 1:5000) antibodies at 4°C overnight, washed three times, and then incubated with 1:5000 Alexafluor-680 or -800 species-specific antibodies for 90 minutes. After three washes, membranes were visualised on a LICOR Odyssey Fc imaging system.

### Northern blotting

Northern blotting was performed as previously described (14). Briefly, A549 cells infected at MOI 3 for 8 hours. Cells were collected, RNA was extracted, then separated by urea-PAGE electrophoresis. RNA was transferred onto nylon membrane overnight then membranes were baked at 68°C for 10 minutes followed by UV crosslinking. Positive control segment 1 and segment 5 transcripts were generated using in vitro transcription assays. Membranes were hybridised overnight with biotinylated probes, then washed, blocked, washed and incubated with HRP-conjugated streptavidin for 30 minutes. Membranes were washed then visualised by exposure to chemiluminescent film.

### Minigenome assay

To measure viral polymerase activity, minigenome reporter assays were performed as described previously (14,41). HEK293T cells were grown to ∼80% confluency in 24-well plates, and then medium was replaced with 400 µl Opti-MEM. Each well was then transfected with 50 ng of PR8 segments 1, 2, 3 and 5 reverse genetics plasmids, or CDLR or CpGH segment 5 plasmids (or no segment 5 plasmid as negative control). In this assay, segment 5 plasmids were titrated, with inputs of 0.005, 0.05, 0.5, 5, 50 or 500 ng, and transfections were performed in technical quadruplicate.

### Serial passage

Wildtype and dinucleotide modified viruses were serially passaged at an MOI of 0.01 for 10 passages in A549 cells and sequenced at passage 10 as previously described (14) with the exception that Seg5 CpGH specific segment 5 primers (amplicons A, B, C and D, **Table 3**) were included in place of the WT PR8 specific segment 5 primers in the pool. Earlier passages were sequenced using set of primers for segment 5 only in 6 overlapping amplicons. These PCRs were performed with the specified primer sets (a, c and e or b, d and f) and Q5 High-Fidelity Polymerase (NEB) according to manufacturer’s instructions. Cycling conditions were 98°C for 30 seconds, then 30 cycles of 98°C for 20 seconds, 55°C for 20 seconds then 72°C for 30 seconds with a final extension of 72°C for 2 minutes. Amplicons purified using the PureLink PCR purification kit (Invitrogen) and subjected to a second PCR round to add partial Illumina sequences and barcodes using the primers in Table 3. Again, reactions were performed using Q5 High-Fidelity Polymerase (NEB) according to manufacturer’s instructions and cycling conditions were 98°C for 30 seconds, then 10 cycles of 98°C for 20 seconds, 55°C for 20 seconds then 72°C for 30 seconds with a final extension of 72°C for 2 minutes. Amplicons were gel purified using the MinElute Gel Extraction kit (Qiagen) and were sent for sequencing using the Amplicon-EZ service (Genewiz). Sequencing data handling and analyses were performed using the Galaxy platform (43). Primer sequences were trimmed from reads using cutadapt and output sequences were joined using fastq-join. Sequences were aligned to the CpGH recoded segments 5 using bowtie2 (44). Variants and coverage levels in the resultant BAM datasets were analysed using iVar variants with tabular outputs.

**Table 3.**
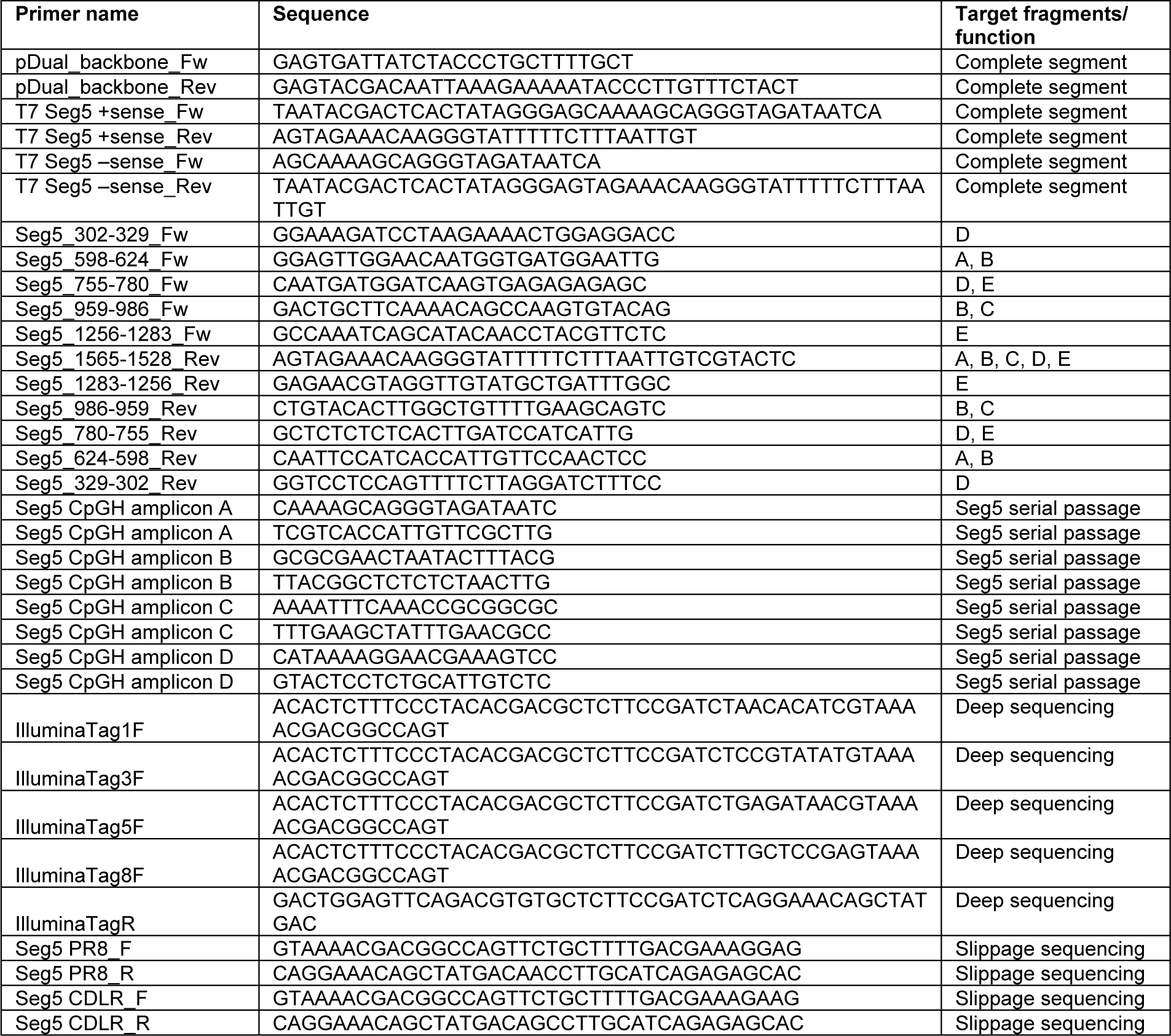
Primers used to generate PCR amplicons.

### Slippage PCR

RNA was extracted from culture supernatants using Viral RNA mini kit (Qiagen) according to manufacturer’s instructions. Extracted RNAs were reverse transcribed using SuperScript III reagents (Invitrogen) with the IAV gRNA specific Uni12 primer (AGCAAAAGCAGG) or cRNA/mRNA specific Uni13 primer (AGTAGAAACAAGG) (45) according to manufacturer’s instructions. PCRs were performed using specific reaction primer pairs specific to the appropriate parental segment (Table 3) and Q5 High-Fidelity Polymerase (NEB) according to manufacturer’s instructions. Cycling conditions were 98°C for 30 seconds, then 30 cycles of 98°C for 20 seconds, 55°C for 20 seconds then 72°C for 30 seconds with a final extension of 72°C for 2 minutes. Amplicons were sent for sequencing using the Amplicon-EZ service (Genewiz) or Sanger sequencing with the reverse primer. Deep amplicon sequencing data handling and analyses were performed using the Galaxy platform (43). Primer sequences were trimmed from reads using cutadapt and output sequences were joined using fastq-join. Sequences were aligned reference genome datasets with 5-15 As in the canonical nucleotide 312-319 region using bowtie2 (44). Coverage levels in the resultant BAM datasets were analysed using Samtools depth. Sanger sequencing traces were visualised using Chromas v2.5.1.

### Serial passage

The virus panel was serially passaged in A549 cells at an MOI of 0.01. Four biological repeats were passaged, with two repeats derived from egg-based virus rescues and two from MDCK-based virus rescues. Viruses were then deep-sequenced after ten passages. For this, RNA was extracted from cell infection supernatants using Qiagen Viral RNA Mini Kit and reverse transcribed using Superscript III (Invitrogen) and primers specific to IAV vRNA (AGCAAAAGCAGG (45)). Whole viral genomes were then amplified in ∼400 base fragments with 60 base overlaps. For PCR, primers from alternate fragments were pooled so that non-overlapping amplicons were generated, with four pools generated in total and segment 5 primers specific to the virus being amplified (Table S1A). PCR reaction mixes included specific primers at a final concentration of 500 nM each and Q5 High-Fidelity Polymerase. Cycling conditions were 98°C for 30 seconds, then 45 cycles of 98°C for 20 seconds, 55°C for 20 seconds, and 72°C for 30 seconds, followed by 72°C final extension for 2 minutes. Multiplexed amplicons were purified using PureLink PCR Purification kit, then DNA was quantified using Qubit dsDNA BR assay kit and pooled in equimolar ratios. Pooled amplicons were sequenced using Genewiz Amplicon-EZ service. Sequencing data was processed and analysed using Galaxy software (43), in which primer sequences were trimmed and then gene fragments were concatenated using PR8 or recoded sequence as the reference genome with Bowtie2 (44). Sequence coverage and variation was analysed using iVar (46).

### Mass spectrometry

To test whether the polyA tract introduced into the CpGH virus resulted in the production of a frame-shifted peptide, ∼ 1 x 10^6^ A549 cells were infected at MOI 3 for 8 hours in the presence or absence of 10 µM MG132 protease inhibitor. Cells were trypsinised and spun down, then stored immediately at -80°C until processing. A549 cells were resuspended in 250 μL 20 mM Tris-pH 7.5, 200 mM NaCl and 1% (w/v) octylthioglucoside then incubated for 1 hour with gentle agitation. Samples were pelleted using centrifugation for 20 minutes at 16000 *g* and 4 °C, and supernatant containing protein was collected. Protein was acetone precipitated then resuspended in 100 mM triethylammonium bicarbonate (TEAB) buffer containing 0.1% SDS, and the protein concentration was measured. 10 μg protein sample was reduced with 5 mM tris(2-carboxyethyl)phosphine (TCEP) for 1 hour at 60 °C then alkylated with 10 mM methylmethanethiosulfonate (MMTS) for 30 minutes in the dark. Due to multiple lysines being encoded at the putative polymerase slippage site, trypsin treatment was unsuitable, and so proteins were instead digested using sequencing grade gluC (Promega Corporation) added to the solution in a 1:20 mass ratio, and incubated at 37 °C. Cleaved peptides were then labelled with iTRAQ reagents (4-Plex system) according to the manufacturer’s instructions (Sciex). iTRAQ labeling regents were dissolved in ethanol, and then transferred to vials containing tryptic peptides (one per sample). Two hours later the reaction was quenched by adding water 1:1. Labelled peptides were pooled and dried under vacuum. Peptides were then fractionated by basic pH reversed-phase chromatography and fractions were desalted on Stage-Tip columns as previously described (47,48). Peptide fractions were loaded on to an Acclaim PepMap100, C_18_, 100 Å, 75 μm × 15 cm column using a Dionex UltiMate RSLCnano System (ThermoFisher Scientific, UK). Peptides were analyzed by a micrOTOF-Q II mass spectrometer (Bruker Daltonics, Bremen, Germany) which using data-dependent acquisition. The *m*/*z* values of tryptic peptides were measured using MS scan (300–2000 *m*/*z*). Raw spectral data were processed using PEAKS software (49) against aligned reference PR8 genome.

### HEK Blue assay

Type I IFN production was measured using HEK Blue reporter assays. Sub-confluent A549 cells in 24-well plates were infected at MOI 10 for 10 hours. Supernatants were then harvested and UV treated to inactivate virus by exposing to 120 mJ/cm^2^ in a UVP CL- 1000 UV crosslinker for 10 minutes. 20 µl UV-treated sample was inoculated onto 4×10^4^ cells/ well of HEK-Blue IFNα/β cells (InvivoGen) in 96-well plates, or titrated human recombinant IFN-β as control (5, 50 or 500 pg/ µl). Cells were incubated at 37°C for 24 hours before supernatants were collected and mixed with QuantiBlue reagent (Invivogen). Colour changes, visible after 15-30 minutes and reflective of the amount of IFN present, were measured by reading absorbance at 620 nm.

### Statistics

All experiments were performed in at least biological triplicate unless otherwise stated, and statistical analyses were only performed on data representative of at least three biological repeats. One way ANOVA tests were performed to assess differences across groups under the same experimental conditions using GraphPad Prism 9.

## ACKNOWLEDGEMENTS

PD, EG and CPS are supported by a BBSRC Institute Strategic Programme grants (BB/P013740/1 and BB/X010937/1). BHT was supported by a Microbiology Society Harry Smith Vacation studentship (GA002550), Wellcome 766 Trust ISSF3 funding and a Carnegie Trust Undergraduate Vacation Scholarship (VAC011866). PD is supported by BBSRC grant no BB/S00114X/1 and the European Union’s Horizon 2020 research and innovation programme under grant agreement no. 727922 (DELTA-FLU). PS is supported by Wellcome Investigator Award 769 (WT103767MA). EG, CPS and BT are also supported by a Wellcome Trust/ Royal Society Sir Henry Dale Fellowship 773 (211222_Z_18_Z). The funders had no role in study design, data collection and analysis, decision to publish, or preparation of the manuscript. For the purpose of open access, the author has applied a CC 775 BY public copyright licence to any Author Accepted Manuscript version arising from this submission.

## REFERENCES

1. Greenbaum, B.D., Levine, A.J., Bhanot, G. and Rabadan, R. (2008) Patterns of evolution and host gene mimicry in influenza and other RNA viruses. PLoS pathogens, 4, e1000079–e1000079.

2. Takata, M.A., Gonçalves-Carneiro, D., Zang, T.M., Soll, S.J., York, A., Blanco-Melo, D. and Bieniasz, P.D. (2017) CG dinucleotide suppression enables antiviral defence targeting non-self RNA. Nature, 550, 124–127.

3. Simmonds, P., Xia, W., Baillie, J.K. and McKinnon, K. (2013) Modelling mutational and selection pressures on dinucleotides in eukaryotic phyla –selection against CpG and UpA in cytoplasmically expressed RNA and in RNA viruses. BMC Genomics, 14.

4. Bird, A.P. (1980) DNA methylation and the frequency of CpG in animal DNA. Nucleic Acids Research, 8, 1499–1504.

5. Cooper, D.N. and Krawczak, M. (1989) Cytosine methylation and the fate of CpG dinucleotides in vertebrate genomes. Human Genetics, 83, 181–188.

6. Antzin-Anduetza, I., Mahiet, C., Granger, L.A., Odendall, C. and Swanson, C.M. (2017) Increasing the CpG dinucleotide abundance in the HIV-1 genomic RNA inhibits viral replication. Retrovirology, 14, 49.

7. Atkinson, N., Witteveldt, J., Evans, D. and Simmonds, P. (2014) The influence of CpG and UpA dinucleotide frequencies on RNA virus replication and characterization of the innate cellular pathways underlying virus attenuation and enhanced replication. Nucleic Acids Research, 42, 4527–4545.

8. Fros, J.J., Dietrich, I., Alshaikhahmed, K., Passchier, T.C., Evans, D.J. and Simmonds, P. (2017) CpG and UpA dinucleotides in both coding and non-coding regions of echovirus 7 inhibit replication initiation post-entry. Elife, 6.

9. Simmonds, P., Tulloch, F., Evans, D.J. and Ryan, M.D. (2015) Attenuation of dengue (and other RNA viruses) with codon pair recoding can be explained by increased CpG/UpA dinucleotide frequencies. Proceedings of the National Academy of Sciences, 112, E3633.

10. Tulloch, F., Atkinson, N.J., Evans, D.J., Ryan, M.D. and Simmonds, P. (2014) RNA virus attenuation by codon pair deoptimisation is an artefact of increases in CpG/UpA dinucleotide frequencies. Elife, 3, e04531.

11. Digard, P., Lee, H.M., Sharp, C., Grey, F. and Gaunt, E. (2020) Intra-genome variability in the dinucleotide composition of SARS-CoV-2. Virus Evol, 6, veaa057.

12. Gaunt, E., Wise, H.M., Zhang, H., Lee, L.N., Atkinson, N.J., Nicol, M.Q., Highton, A.J., Klenerman, P., Beard, P.M., Dutia, B.M. et al. (2016) Elevation of CpG frequencies in influenza A genome attenuates pathogenicity but enhances host response to infection. Elife, 5, e12735–e12735.

13. Gaunt, E.R. and Digard, P. (2021) Compositional biases in RNA viruses: Causes, consequences and applications. *WIREs RNA*, e1679.

14. Sharp, C.P., Thompson, B.H., Nash, T.J., Diebold, O., Pinto, R.M., Thorley, L., Lin, Y.-T., Sives, S., Wise, H., Clohisey Hendry, S. et al. (2023) CpG dinucleotide enrichment in the influenza A virus genome as a live attenuated vaccine development strategy. PLOS Pathogens, 19, e1011357.

15. Svitkin, Y.V., Cammack, N., Minor, P.D. and Almond, J.W. (1990) Translation deficiency of the sabin type 3 poliovirus genome: Association with an attenuating mutation C472 → U. Virology, 175, 103–109.

16. Eisfeld, A.J., Neumann, G. and Kawaoka, Y. (2015) At the centre: influenza A virus ribonucleoproteins. Nature reviews. Microbiology, 13, 28–41.

17. Te Velthuis, A.J. and Fodor, E. (2016) Influenza virus RNA polymerase: insights into the mechanisms of viral RNA synthesis. Nature reviews. Microbiology, 14, 479–493.

18. Poon, L.L.M., Pritlove, D.C., Fodor, E. and Brownlee, G.G. (1999) Direct Evidence that the Poly(A) Tail of Influenza A Virus mRNA Is Synthesized by Reiterative Copying of a U Track in the Virion RNA Template. 73, 3473–3476.

19. García-Sastre, A., Egorov, A., Matassov, D., Brandt, S., Levy, D.E., Durbin, J.E., Palese, P. and Muster, T. (1998) Influenza A Virus Lacking the NS1 Gene Replicates in Interferon-Deficient Systems. Virology, 252, 324–330.

20. Phan, T., Fay, E.J., Lee, Z., Aron, S., Hu, W.-S. and Langlois, R.A. (2021) Segment-Specific Kinetics of mRNA, cRNA, and vRNA Accumulation during Influenza Virus Infection. 95, e02102–02120.

21. Michalak, P., Soszynska-Jozwiak, M., Biala, E., Moss, W.N., Kesy, J., Szutkowska, B., Lenartowicz, E., Kierzek, R. and Kierzek, E. (2019) Secondary structure of the segment 5 genomic RNA of influenza A virus and its application for designing antisense oligonucleotides. Sci Rep, 9, 3801.

22. Luo, X., Wang, X., Gao, Y., Zhu, J., Liu, S., Gao, G. and Gao, P. (2020) Molecular Mechanism of RNA Recognition by Zinc-Finger Antiviral Protein. Cell Reports, 30, 46–52.e44.

23. Li, M.M.H., Lau, Z., Cheung, P., Aguilar, E.G., Schneider, W.M., Bozzacco, L., Molina, H., Buehler, E., Takaoka, A., Rice, C.M. et al. (2017) TRIM25 enhances the antiviral action of zinc-finger antiviral protein (ZAP). PLoS Pathogens, 13, e1006145.

24. Zheng, X., Wang, X., Tu, F., Wang, Q., Fan, Z. and Gao, G. (2017) TRIM25 is required for the antiviral activity of zinc finger antiviral protein. Journal of Virology, 91, e00088–00017.

25. Choudhury, N.R., Heikel, G., Trubitsyna, M., Kubik, P., Nowak, J.S., Webb, S., Granneman, S., Spanos, C., Rappsilber, J., Castello, A. et al. (2017) RNA-binding activity of TRIM25 is mediated by its PRY/SPRY domain and is required for ubiquitination. BMC Biology, 15, 105.

26. Choudhury, N.R., Trus, I., Heikel, G., Wolczyk, M., Szymanski, J., Bolembach, A., Dos Santos Pinto, R.M., Smith, N., Trubitsyna, M., Gaunt, E., et al. (2022) TRIM25 inhibits influenza A virus infection, destabilizes viral mRNA, but is redundant for activating the RIG-I pathway. Nucleic Acids Research, 50, 7097–7114.

27. Arevalo, C.P., Bolton, M.J., Le Sage, V., Ye, N., Furey, C., Muramatsu, H., Alameh, M.-G., Pardi, N., Drapeau, E.M., Parkhouse, K. et al. (2022) A multivalent nucleoside-modified mRNA vaccine against all known influenza virus subtypes. 378, 899–904.

28. Singanayagam, A., Zambon, M., Lalvani, A. and Barclay, W. (2018) Urgent challenges in implementing live attenuated influenza vaccine. The Lancet Infectious Diseases, 18, e25–e32.

29. Chen, J.-R., Liu, Y.-M., Tseng, Y.-C. and Ma, C. (2020) Better influenza vaccines: an industry perspective. Journal of Biomedical Science, 27.

30. Jang, Y.H. and Seong, B.-L. (2012) Principles underlying rational design of live attenuated influenza vaccines. Clin Exp Vaccine Res, 1, 35–49.

31. Kassianos, G., MacDonald, P., Aloysius, I. and Reynolds, A. (2020) Implementation of the United Kingdom’s childhood influenza national vaccination programme: A review of clinical impact and lessons learned over six influenza seasons. Vaccine, 38, 5747–5758.

32. Maassab, H.F. and DeBorde, D.C. (1985) Development and characterization of cold-adapted viruses for use as live virus vaccines. Vaccine, 3, 355–369.

33. Moratorio, G., Henningsson, R., Barbezange, C., Carrau, L., Bordería, A.V., Blanc, H., Beaucourt, S., Poirier, E.Z., Vallet, T., Boussier, J. et al. (2017) Attenuation of RNA viruses by redirecting their evolution in sequence space. Nature Microbiology, 2, 17088.

34. Mueller, S., Coleman, J.R., Papamichail, D., Ward, C.B., Nimnual, A., Futcher, B., Skiena, S. and Wimmer, E. (2010) Live attenuated influenza virus vaccines by computer-aided rational design. Nature Biotechnology, 28, 723–726.

35. Yang, C., Skiena, S., Futcher, B., Mueller, S. and Wimmer, E. (2013) Deliberate reduction of hemagglutinin and neuraminidase expression of influenza virus leads to an ultraprotective live vaccine in mice. Proc Natl Acad Sci U S A, 110, 9481–9486.

36. Corbett, K.S., Edwards, D.K., Leist, S.R., Abiona, O.M., Boyoglu-Barnum, S., Gillespie, R.A., Himansu, S., Schäfer, A., Ziwawo, C.T., DiPiazza, A.T. et al. (2020) SARS-CoV-2 mRNA vaccine design enabled by prototype pathogen preparedness. Nature, 586, 567–571.

37. Shaw, A.E., Rihn, S.J., Mollentze, N., Wickenhagen, A., Stewart, D.G., Orton, R.J., Kuchi, S., Bakshi, S., Collados, M.R., Turnbull, M.L. et al. (2021) The antiviral state has shaped the CpG composition of the vertebrate interferome to avoid self-targeting. PLoS biology, 19, e3001352.

38. Ficarelli, M., Wilson, H., Pedro Galão, R., Mazzon, M., Antzin-Anduetza, I., Marsh, M., Neil, S.J.D. and Swanson, C.M. (2019) KHNYN is essential for the zinc finger antiviral protein (ZAP) to restrict HIV-1 containing clustered CpG dinucleotides. Elife, 8, e46767.

39. de Wit, E., Spronken, M.I., Bestebroer, T.M., Rimmelzwaan, G.F., Osterhaus, A.D. and Fouchier, R.A. (2004) Efficient generation and growth of influenza virus A/PR/8/34 from eight cDNA fragments. Virus Res, 103, 155–161.

40. Hutchinson E. C., Curran M. D., Read E. K., Gog J. R. and Digard, P. (2008) Mutational Analysis of cis-Acting RNA Signals in Segment 7 of Influenza A Virus. Journal of Virology, 82, 11869–11879.

41. Wise HM., Foeglein, A., Sun, J., Dalton Rosa, M., Patel, S., Howard, W., Anderson E. C., Barclay W. S. and Digard, P. (2009) A Complicated Message: Identification of a Novel PB1- Related Protein Translated from Influenza A Virus Segment 2 mRNA. Journal of Virology, 83, 8021–8031.

42. Carrasco, M., Amorim, M.J. and Digard, P. (2004) Lipid raft-dependent targeting of the influenza A virus nucleoprotein to the apical plasma membrane. *Traffic (Copenhagen*, Denmark), 5, 979–992.

43. Afgan, E., Baker, D., Batut, B., van den Beek, M., Bouvier, D., Čech, M., Chilton, J., Clements, D., Coraor, N., Grüning, B.A., et al. (2018) The Galaxy platform for accessible, reproducible and collaborative biomedical analyses: 2018 update. Nucleic Acids Research, 46, W537–W544.

44. Langmead, B., Trapnell, C., Pop, M. and Salzberg, S.L. (2009) Ultrafast and memory-efficient alignment of short DNA sequences to the human genome. Genome Biology, 10, R25.

45. Hoffmann, E., Stech, J., Guan, Y., Webster, R.G. and Perez, D.R. (2001) Universal primer set for the full-length amplification of all influenza A viruses. Arch Virol, 146, 2275–2289.

46. Grubaugh, N.D., Gangavarapu, K., Quick, J., Matteson, N.L., De Jesus, J.G., Main, B.J., Tan, A.L., Paul, L.M., Brackney, D.E., Grewal, S. et al. (2019) An amplicon-based sequencing framework for accurately measuring intrahost virus diversity using PrimalSeq and iVar. Genome Biology, 20, 8.

47. Rappsilber, J., Mann, M. and Ishihama, Y. (2007) Protocol for micro-purification, enrichment, pre-fractionation and storage of peptides for proteomics using StageTips. Nature Protocols, 2, 1896–1906.

48. Withatanung, P., Kurian, D., Tangjittipokin, W., Plengvidhya, N., Titball, R.W., Korbsrisate, S. and Stevens, J.M. (2019) Quantitative Proteomics Reveals Differences in the Response of Neutrophils Isolated from Healthy or Diabetic Subjects to Infection with Capsule-Variant Burkholderia thailandensis. Journal of Proteome Research, 18, 2848–2858.

49. Zhang, J., Xin, L., Shan, B., Chen, W., Xie, M., Yuen, D., Zhang, W., Zhang, Z., Lajoie, G.A. and Ma, B. (2012) PEAKS DB: de novo sequencing assisted database search for sensitive and accurate peptide identification. Molecular & cellular proteomics : MCP, 11, M111.010587.

